# HIVtat Alters Epithelial Differentiation State and Increases HPV16 Infectivity in Oral Keratinocytes

**DOI:** 10.1101/2023.03.08.531567

**Authors:** William T Seaman, Victoria Madden, Jennifer Webster-Cyriaque

## Abstract

Human Papillomavirus (HPV)-associated oral disease has increased during the era of HIV antiretroviral therapy. HPV and HIV proteins may be co-present at mucosal surfaces. Recent published studies have determined that HIVtat is secreted in the saliva and has been detected in oral mucosa even in the context of antiretroviral therapy. We hypothesized that HIVtat promoted oral HPV pathogenesis. Clinical HPV16 cloned episomes were introduced into differentiated oral epithelial cells (OKF6tert1). HIVtat mediated transactivation, DNA damage, oxidative stress, and effects on cellular differentiation were assessed. Detection of keratin 10 and of loricrin confirmed terminal differentiation. Sodium butyrate-treated (NaB) cells demonstrated an eight-fold increase in cross-linked involucrin, suggesting full terminal differentiation. HIVtat modulated this differentiation both in the presence and absence of NaB. Later viral events, including E6* and E1^E4 gene expression were assessed. HIVtat mediated relief of repressed L1 expression that mapped to a known inhibitory region (nucleotides 5561-6820). Viruses from HIVtat co-expressing cells exhibited robust de novo HPV16 infection. In conclusion, a novel oral keratinocyte monolayer system supported replication of an HPV16 clinical isolate where direct HIVtat and oral HPV interactions enhanced HPV de novo infection.

## Introduction

The link between HPV and cervical cancer has been well established (7, 41). Less information exists for the role HPV plays in the establishment of oral cancers. Recent data from our lab and others has shown that HPV is associated with at least 15-20% of oral cancers (2, 15, 23, 39, 42, 45). High-risk types of HPV DNA have been previously consistently detected in 20% of head and neck squamous cell carcinomas (HNSCC) overall with most originating in the oropharynx. More recently, HPV has been detected in up to 65% of oral cancers and has been detected at a 1.5-4.0 fold higher rate in HIV positive subjects (5, 49).

Oral HPV associated disease is even more striking in the context of immune suppression. The prevalence of oral HPV in HIV infected persons (26%) is 3.5 times higher than the US general population (7%) (5). Recent studies show higher risk of oropharyngeal squamous cell carcinoma (SCC) in HIV-infected patients compared with general population (6, 11). HPV associated disease persists in the era of highly active antiretroviral therapy (1, 10, 22, 36). Higher proportion of individuals with HPV associated oral warts found among those on HAART (23%) than among those not taking HAART (5%) (26). Oral warts were found to be associated ≥1 log10 decrease in HIV-1 RNA (30). While antiretroviral therapy does decrease HIV viral loads, it does not completely block the secretion of HIVtat protein (32). Although HIV infects lymphocytes, HIVtat has been detected by immunopreciptation in saliva of 33% HIV positive individuals (48). HIVtat is a potent transcription factor known to enhance cellular DNA damage and oxidative stress (38, 46). HIVtat is easily taken up by multiple cell types including epithelial cells (20, 25). Further, it has been shown that salivary HIVtat, at physiologic concentrations, can penetrate oral epithelial cultures (48). It has recently been demonstrated that CD68 and CD3 positive immune cells trafficking through the oral mucosa of HIV positive individuals express both HIVtat and gp120. The expression of HIVtat is reduced by at least 50% in those individuals on HAART, however, HIVtat remains detectable in the oral epithelium and its associated immune cell complement (48).

In order to better define steps important to the development of oral tumors, it is critical to understand HPV infection of oral epithelial cells in the context of HIV. A simple culture system was developed utilizing full length wild type oral HPV clinical isolates and immortalized oral keratinocytes to investigate the HPV lifecycle and factors important to HPV replication including DNA damage and terminal differentiation. The culture system facilitated the definition of viral and cellular factors critical to the spread of HPV in the oral cavity. LoxP-containing HPV clones from oral clinical samples were generated by PCR amplifying entire HPV genomes with type-specific primers that included loxP sequences. Transfected cells expressed early and late viral mRNAs and effected cellular responses similar to other culture systems. Using this culture system, HPV viral particles with packaged episomes were produced indicating that these cells were capable of supporting the full viral life cycle. The use of these oral keratinocytes provided a straight forward infection model in which to determine the role of HIVtat in oral HPV infections. HIVtat and HPV interactions in oral epithelia were investigated. The above described system has provided important insights to the relationship between HPV and HIV within oral epithelia. Interestingly, the expression of HIVtat in HPV expressing oral epithelial cells resulted in modifications in both cellular differentiation state and in HPV gene expression. Further, de novo HPV infection was enhanced in the presence of HIVtat.

## Materials and Methods

### Cell culture

OKF6tert1 cells are a telomerase immortalized oral keratinocyte cell line and were provided by James Rheinwald and propagated as described (18). Briefly, OKF6tert1 cells were grown in Keratinocyte-SF (KSF) media (Gibco) supplemented with 0.2 ng/ml EGF and 25 μg/ml BPE. CaCl_2_ was added to a final concentration of 0.4 mM. Caski cells were grown in Dulbecco’s modified Eagles medium supplemented with 10% Fetal Calf serum. All cells were maintained in a humidified incubator at 37° C with 5% CO_2_. Cells were transfected with Fugene-6 (Promega) according to the manufacturer’s instructions. Prior to treatment with NaB or CaCl2 cells were washed once with PBS and overlaid with fresh KSF media without supplements. After 24 hours, media was removed and replaced with KSF media without supplements containing either 0.6 mM NaB or 2.0 mM CaCl2.

### Whole genome amplification

Isolation of genomic DNA from oral cancer biopsy has been previously described (42). Whole genome amplification of HPV16 from oral cancer DNA was performed with an Expand High Fidelity PCR system (Roche) according to the manufacturer’s instructions using HPV type 16-specific primers, 16loxpF and 16loxpR. One nanogram of genomic DNA was used in the reaction. Thermal cycle conditions for amplification were as follows: 94°C 2 minute initial denaturation followed by 10 cycles of 94°C for 15 seconds (denature), 60°C for 30 seconds (primer annealing) and 68°C for 7 minutes (extension). These 10 cycles were followed 20 cycles of the same thermal cycle profile except that 15 seconds were added to each successive 7 minute extension at 68°C. PCR products were analyzed by agarose gel electrophoresis.

### Plasmids and primers

The Cre expression plasmid, pCre-CMVPro, was a gift from the UNC Gene Therapy Core Facility. The HIV LTR luciferase reporter plasmid, pHIVLTRluc, was a gift from Lishan Su. Primer sequences used for PCR amplification and shown in Table S1. All primers were manufactured at the UNC Nucleic Acid Core Facility. HPV16 whole genome PCR product was cloned into the HindIII/XmaI sites of pcDNA3 (Invitrogen) to construct pCMVHPV16neo. HPV16 E6 and E7 expression plasmids, pcDNAHPV16E6 and pcDNAHPV16E7 were constructed by PCR amplification of either the E6 coding region (83–559) with HPV16E6F/R primers or the E7 coding region (562–858) with HPV16E7F/R primers using Caski cell DNA as a template followed by cloning into the HindIII/XhoI sites of pcDNA3. HPV16 E2 expression plasmid, pcDNAHPV16E2 was constructed by PCR amplification of the E2 coding region (2755–3852) with HPV16E2F/R primers using Caski cell DNA as a template followed by cloning into the HindIII/XhoI sites of pcDNA3. HPV16 L1 expression plasmid, pcDNAHPV16L1 was constructed by PCR amplification of the L1 coding region (5561–7156) with HPV16L1F/R primers using pCMVHPV16neo as a template followed by cloning into the EcoRV/XhoI sites of pcDNA3.The HPV LCR luciferase reporter plasmid, pHPV16luc was constructed by inserting the HPV16 LCR (7009-7905/1-84), PCR amplified with HPV16LCRF/R primers, into the MluI/BglII sites of pGL2-Basic (Promega). The following reagent was obtained through the NIH AIDS Reagent Program, Division of AIDS, NIAID, NIH: pTatC6H-1 from Dr. Abhay Patki and Dr. Michael Lederman (37). The HIVtat coding region was by PCR amplified from pTatC6H-1 using HIVtatFF and HIVtat3’RR primers followed by digestion with EcoRI. To make pCMV-tat, the PCR fragment was inserted into pCMV-HA that had been digested with StuI and EcoRI. This removed the HA tag from the resulting construct. The GFP expression plasmid, pcDNAGFP was constructed by inserting the GFP coding region PCR amplified with GFPF/R primers into the HindIII/KpnI site of pcDNA3. The KpnI/XhoI fragment of pcDNA3HPV16L1 (5561–7156) was inserted into the KpnI/XhoI sites of pcDNAGFP to produce pGFPL1. The KpnI/BamHI fragment of pcDNA3HPV16L1 (5561–6152) was inserted into the KpnI/BamHI sites of pcDNAGFP to produce pGFPL1BamHI5’. The KpnI/EcoRI fragment of pcDNA3HPV16L1 (5561–6820) was inserted into the KpnI/EcoRI sites of pcDNAGFP to produce pGFPL1EcoRI5’. The BamHI/XhoI fragment of pcDNA3HPV16L1 (6152–7156) was inserted into the BamHI/XhoI sites of pcDNAGFP to produce pGFPL1BamHI3’. The EcoRI/XhoI fragment of pcDNA3HPV16L1 (6820–7156) was inserted into the EcoRI/XhoI sites of pcDNAGFP to produce pGFPL1EcoRI3’. Primer sequences for PCR amplification are shown in Supplemental Table 1. All plasmid constructs were verified by sequencing.

### RT-qPCR

Total RNA was isolated from transfected cells using Trizol according to the manufacturer’s instructions (Invitrogen). Briefly, cells were washed twice with 1X PBS before the addition of Trizol. Total RNA was incubated with RQ1 RNase-free DNase (Promega) for 45 minutes at 37° C prior use in reverse transcription reactions. After incubation DNase activity was destroyed by heating to 70° C for 10 minutes. 200 ng of total RNA was reverse transcribed using SuperScript III (Invitrogen) and random primers. Reactions were performed at 50° C for 1 hour. Following incubation, reactions were diluted to 100 μl with water and 2 μl were used for qPCR. Real time PCR reactions were performed in triplicate using Roche Lightcycler Sybr Green master mix. Each reaction consisted of 1X Roche Lightcycler Sybr Green master mix and gene-specific primers (Supplemental Table 1) in a 10 μl reaction. Thermal cycle conditions consisted of an initial denaturation incubation at 95° C for 10 minutes followed by 45 cycles of alternating 95° C incubations for 10 seconds, 60° C incubations for 10 seconds and 72° C incubations for 30 seconds. Fluorescence was detected after every 72° C extension incubation. GAPDH levels were used to normalize experimental gene transcript levels.

### Western blot analysis

Total protein was isolated from cells using Trizol (Invitrogen) according to the manufacturer’s instructions. Final protein pellets were resuspended in 1X sample buffer (250 mM Tris, 8.0, 2% SDS, 20% glycerol, 50 mM DTT). Pellets in buffer were incubated at 65° C until dissolved. Twenty-five micrograms of total protein were loaded onto precast NuPage 4-12% Bis-Tris gels (Invitrogen). Proteins were electroblotted to nitrocellulose paper using western blot transfer buffer (25 mM Tris base, 192 mM glycine, 20% methanol). Following transfer, blots were blocked by incubation with 5% nonfat dry milk in 1X PBS containing 0.1% Tween-20 (1X PBS-T) for 1 hour at room temperature. Following blocking, blots were incubated with primary antibody for 1 hour at room temperature. Blots were washed twice with 1X PBS-T followed by incubation for 1 hour with HRP-conjugated secondary antibody (Promega) diluted 1:10,000 in PBS-T, 5% milk. Blots were washed twice with PBS-T and protein bands were detected by ECL after incubation with SuperSignal for 5 minutes according to the manufacturer’s instructions (ThermoScientific). The following primary antibodies were used to detect proteins on western blots: anti-involucrin (sc-28557; Santa Cruz), anti-loricrin (PRB-145P; Biolegend), anti-GAPDH (sc-25778; Santa Cruz), anti-γ-H2A.X Ser 139 (sc-101696; Santa Cruz), anti-p-Chk2 Thr 68 (sc-16297-R; Santa Cruz), anti-actin (sc-1616-R; Santa Cruz), anti-p38 MAPK (9212S; Cell Signaling), anti-phospho p38MAPK (9215S; Cell Signaling), anti-p53 (sc-6243; Santa Cruz), anti-histone H3 (D1H2; Cell Signaling), The following reagent was obtained through the NIH AIDS Reagent Program, Division of AIDS, NIAID, NIH: Antiserum to HIV-1 Tat from Dr. Bryan Cullen (catalog #705) (27).

### Quantitation of DNase-resistant viral DNA

Media from transfected cells were passed through a 0.45 micron filter. 200 ml of media was treated with 4 U of RQ1 RNase-free DNase (Promega) for 1 hour at 37°C. Following incubation DNase activity was destroyed by heating to 70° C for 15 minutes. DNA was isolated with a Qiagen DNeasy Blood & Tissue kit according to the manufacturer’s instructions. HPV16 DNA was detected by qPCR using 1X Roche Lightcycler Sybr Green master mix and HPV16 E7-specific primers. Thermal cycle conditions consisted of an initial denaturation incubation at 95° C for 10 minutes followed by 45 cycles of alternating 95° C incubations for 10 seconds, 60° C incubations for 10 seconds and 72° C incubations for 30 seconds. Fluorescence was detected after every 72° C extension incubation.

### Negative staining TEM

HPV particles were detected by TEM as previously described (44). Briefly, media from transfected cells was centrifuged for 20,000 X G for 30 minutes to remove cellular debris followed by centrifugation at 100,000 X G for 1 hour to pellet viral particles. Pellets were resuspended in 50 μl of water and fixed with 3% glutaraldehyde. Particles were stained with uranyl acetate and analyzed by TEM.

### Immunofluorescence Assay (IFA)

Following transfection, cells were washed with 1X PBS and fixed with methanol/acetone. Blocking was performed for 1 hour with 1X PBS containing 5% normal goat serum. For de novo infection, filtered media from transfected cells was used as an inoculum for naïve OKF6tert1. 96 hours post infection cells were fixed with methanol/acetone. HPV16 E7 was detected using mouse anti-HPV16 E7 (sc-6981; Santa Cruz Biotech) and HPV L1 was detected with CAMVIR-1 antibody (sc-47699; Santa Cruz Biotech). Fixed cells were overlaid with 1:500 dilution of primary antibody and incubated overnight at 4° C. Cells were washed three time with 1X PBS (5 minutes/wash) followed by overlaying with 1:500 dilution of Alexa Fluor 488 goat-anti mouse IgG (A28175; Pierce) for 1 hour at room temperature. Rabbit anti-HIVtat antiserum was obtained through the NIH AIDS Reagent Program (cat. #705), Division of AIDS, NIAID, NIH: Antiserum to HIV-1 Tat from Dr. Bryan Cullen (27). Bound anti-HIVtat antibody was detected with Alexa Fluor 594 goat anti-rabbit IgG secondary antibody (A11072; ThermoFisher). To stain nuclei, cells were overlaid with DAPI for 5 minutes followed by washing three time with 1X PBS (5 minutes/wash). Fluorescence was detected with an Olympus IX81 fluorescent microscope.

## Results

### HIVtat was active in oral epithelial cells but did not activate the HPV LCR

Given the increase in HPV-associated disease within the context of HIV and the biochemical and biophysical properties of HIVtat, the impact of HIVtat on the HPV16 life cycle was assessed within the HPV16/OKF6tert1 system. HIVtat expression in OKF6tert cells was confirmed by the detection of the luciferase produced from an HIVLTR luciferase reporter plasmid after cotransfection with a HIVtat expression vector in the presence and absence of NaB (Figure 1A). HPV gene expression in the presence of HIVtat was confirmed in these cells by the diminished detection of p53 in the HPV expressing cells, suggesting functional HPV E6 protein was present (Figure 1B, lanes 5-8). HIVtat had been previously shown to enhance the activity of the HPV LCR in HeLa cells (47). In this study, while HIVtat effectively transactivated the HIVLTR demonstrating a 50 fold increase in luciferase reporter activity (Figure 1C right panel), HIVtat did not increase reporter activity from the HPV LCR in OKF6tert1 cells (Figure 1C left panel).

**Figure 1.**
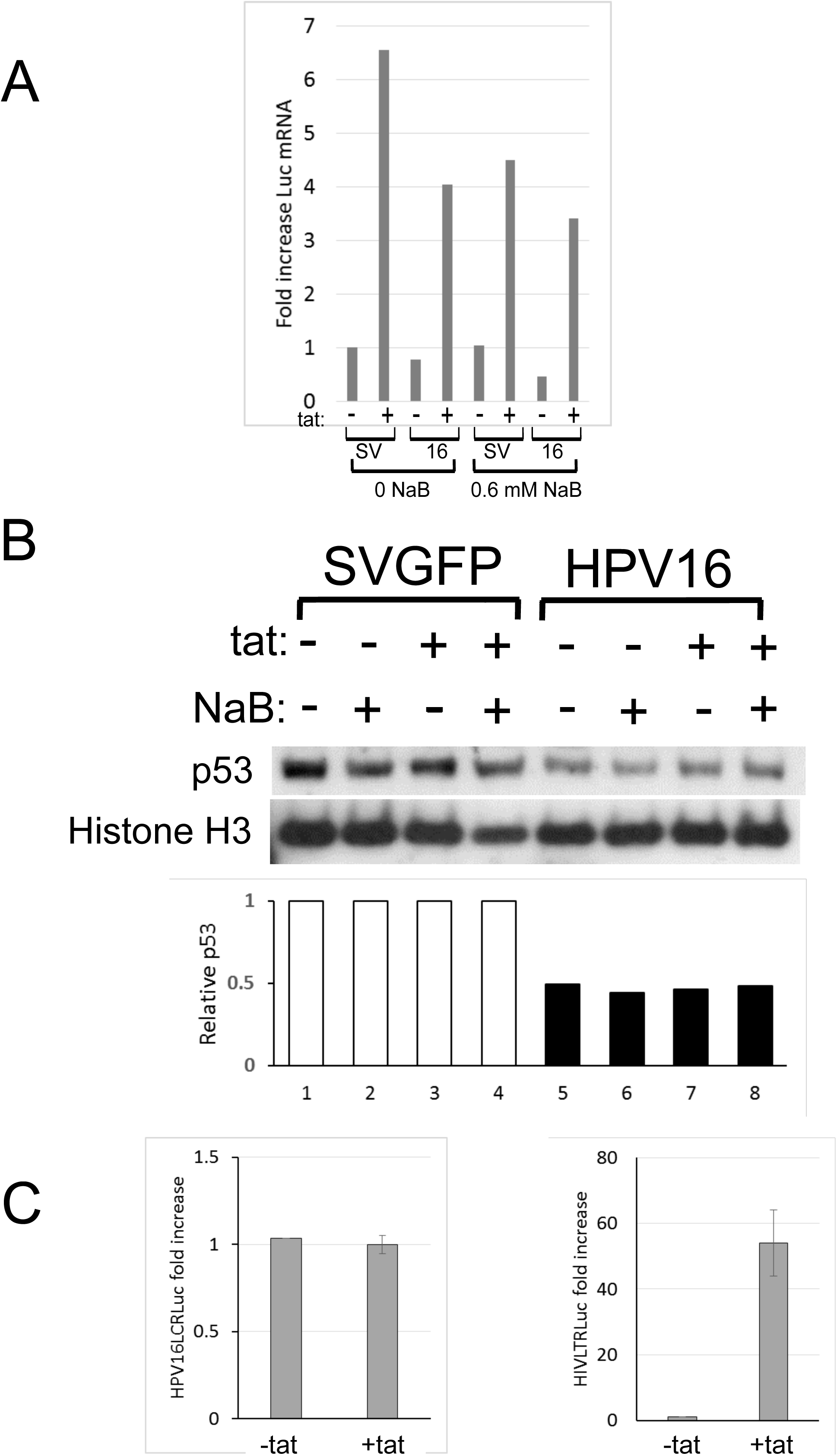
p53 is downregulated in OKF6tert1cells harboring HPV16 episomes. A. The level of luciferase gene expression driven by the HIV LTR was measured by RT-qPCR using Luciferase-specific primers to ensure that HIVtat was being expressed in these cells. Fold increase was determined after normalization to GAPDH levels. B. Western blot analysis was performed on total cell extracts obtained after introduction of HPV16 episomes into OKF6tert1 cells in the presence or absence of HIVtat followed by treatment with 0.6 mM NaB. The blot was probed with anti-p53 antibody to determine p53 levels. Total histone H3 was used as loading controls. Relative p53 levels are indicated after normalization to the total H3 signal. **C.** HIVtat did not significantly affect transactivation of the HPV16 LCR in OKF6tert1 cells. OKF6tert1 cells were cotransfected with an HPV16 LCR/luciferase reporter plasmid and either empty vector or a tat-expression vector (pCMV-tat). No increase in luciferase expression from the HPV16LCR construct was detected when HIVtat was included (left panel). HIVtat was capable of significantly increasing HIVLTR transactivation after cotransfection of OKF6tert1 cells with HIVLTR/luciferase plasmids and the HIVtat expression vector (right panel).

### HIVtat does not enhance DNA damage in HPV expressing cells but did enhance epithelial differentiation state

Others have shown that activation of the DNA damage response is favorable to HPV replication (33). The ability of HIVtat to induce cellular responses, like DNA damage, that are favorable to virus production suggested that HIVtat could affect HPV16 replication. A TUNEL assay revealed DNA damage induced by expression of HIVtat in OKF6tert1 cells, independent of HPV presence (Figure 2A). The increase in phospho-chk2 associated with the expression of HIVtat in OKF6tert1 cells confirmed the DNA-damage response in the absence of HPV (Figure 2B (left panel) lanes 1 and 2). HPV16 expressing OKF6tert1 cells demonstrated expression of phopho-chk-2 compared to the vector control (Figure 2B (left panel) lanes 1 and 3). In the presence of HIVtat, HPV 16 transfected cells did not reveal enhanced expression of phopho-chk-2 (Figure 2B (left panel) lanes 3 and 4). HIVtat did not increase in γ-H2AX, another marker of DNA damage, in the absence of HPV (Figure 2B (right panel), compare lanes 1 and 5). Similarly, coexpression of HIVtat and HPV16 oncogenes E6 and E7, did not result in a synergistic effect on the DNA damage response (Figure 2B (right panel) lanes 2-4 compared to lanes 6-8). HIVtat-induced DNA damage could be the result of increased oxidative stress (51). To determine if HIVtat increased oxidative stress in OKF6tert1 cells, the levels of glutathione peroxidase (GPX) and super oxide dismutatase (SOD) transcripts were determined by RT-qPCR (Figure 2C). While the oxidative stress response was not directly assessed, decreased expression of oxidative stress response genes occurred in cells expressing HIVtat compared with vector control. Statistically significant decreases in GPX expression were not detected (p=0.18), however, a modest but statistically significant decrease in SOD gene expression was detected (p=0.03) (Figure 2C). As a control, HIV LTR responsiveness was assessed with each of these constructs as well (p=0.02).

**Figure 2.**
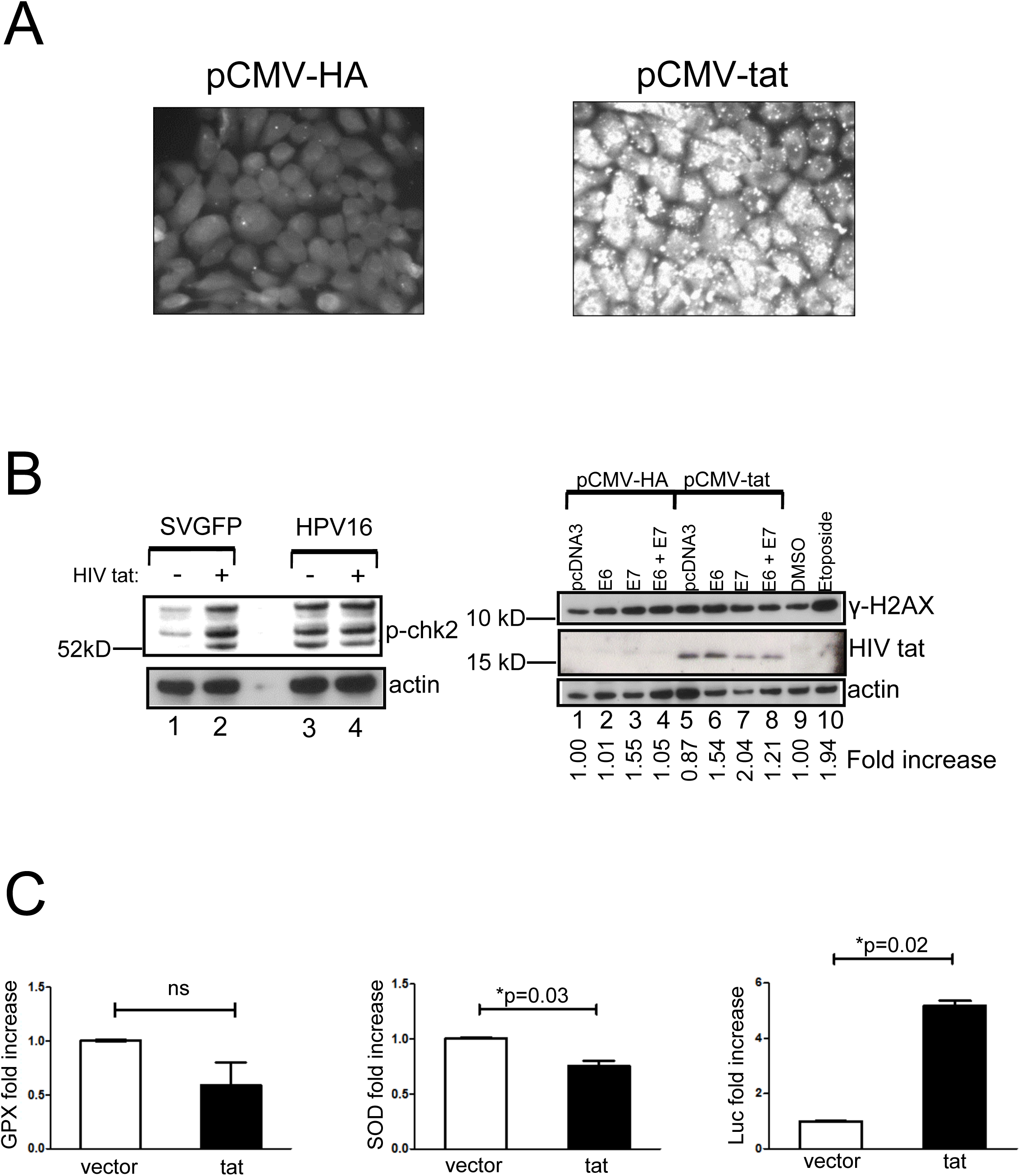
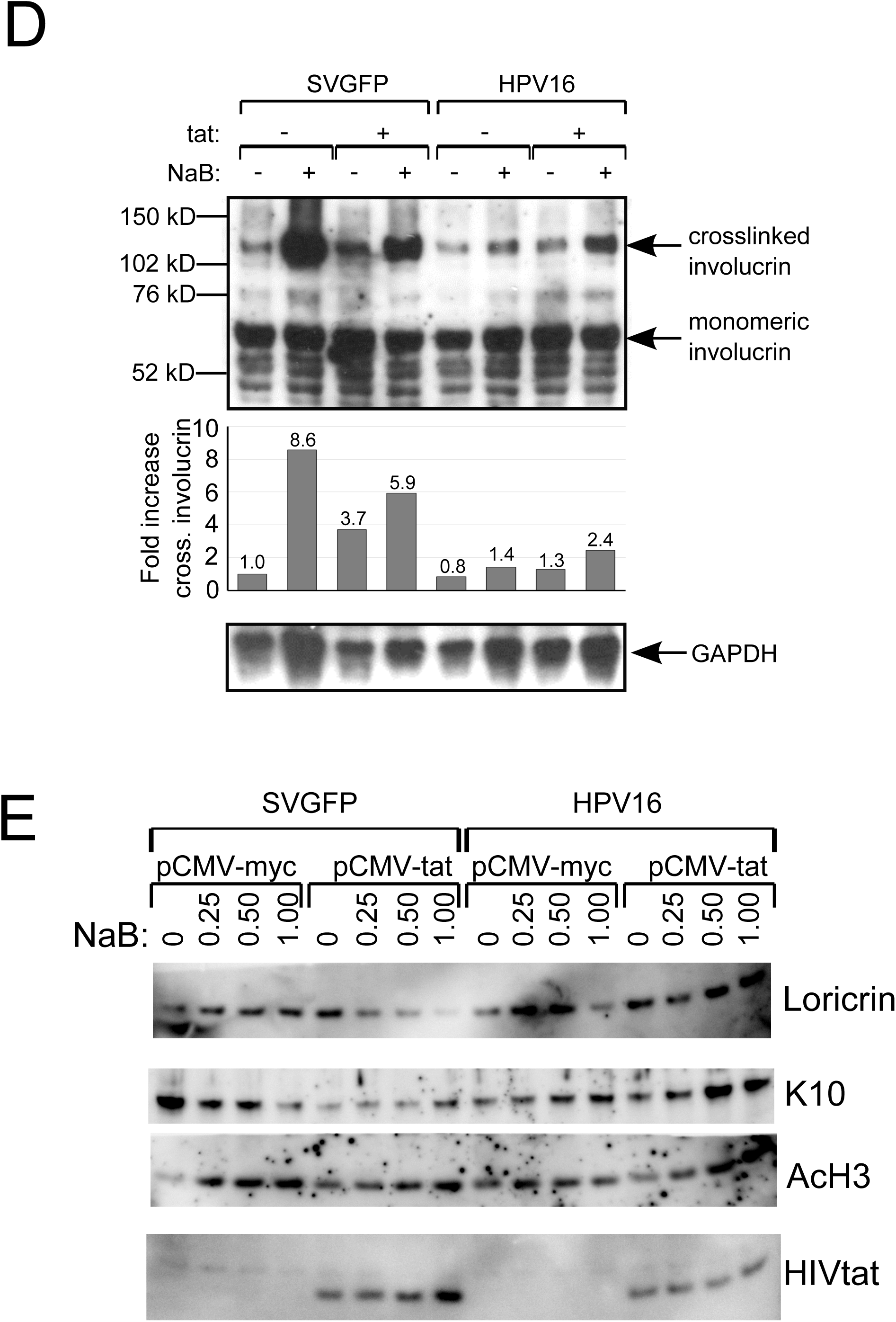
Modulation of cellular responses in the presence of HIVtat and HPV 16 episomes. A DNA damage response was induced and differentiation was suppressed in OKF6tert1 cells expressing HIVtat and harboring HPV16 episomes. **A.** DNA damage occurs in OKF6tert1 cells expressing HIVtat. Cells were transfected with control vector (pCMV-HA) or HIVtat expression vector (pCMV-tat). 48 hours post transfection cells were assayed for DNA damage by TUNEL assay. **B.** The levels of phospho-chk2 (p-chk2) were analyzed in either control cells or HPV16+ cells in the presence or absence of HIVtat. OKF6tert1 cells were cotransfected with pSVGFP and either control vector (pCMV-HA) or the HIVtat expression vector, pCMV-tat. Another set of cells were cotransfected with pHPV16loxp and either control vector (pCMV-HA) or the HIVtat expression vector, pCMV-tat. 48 hours post transfection cell lysates were obtained and analyzed by western blot analysis for the DNA damage marker, p-chk2. β-actin was detected on the same blot to show equal loading of the lysates (left panel). OKF6tert1 cells expressing HPV16 E7 and HIVtat have an increased DNA damage response. OKF6tert1 cells were cotransfected with either HPV16 E6 or E7 expression vectors and either control vector (pCMV-HA) or the HIV-tat expression vector, pCMV-tat. 48 hours post transfection cell lysates were obtained and analyzed by western blotting for the DNA damage marker, γ-H2AX. HIVtat was detected using a rabbit anti-tat antibody. Cell lysate from OKF6tert1 cells treated with the DNA damage-inducing agent, etoposide was used as a positive control. Numbers below each lane indicate the fold increase in γ-H2AX over control transfected cells (pcDNA3 + pCMV-HA) as determined by densitometry after normalization to β-actin (right panel). **C.** OKF6tert1 cells were transfected with either control vector (pCMV-HA) or the tat expression vector, pCMV-tat. 48 hours post transfection total RNA was isolated and used to perform RT-qPCR for the detection of transcripts encoding the oxidative stress markers, glutathione peroxidase (GPX) and superoxide dismutase (SOD). While HIVtat expression caused a slight decrease in GPX transcripts it was not statistically significant (p=0.18). However, HIVtat expression did result in a statistically significant decrease in SOD transcripts (p=.0019). **D.** Cell lysates from OKF6tert1 cells treated with vehicle or 0.6mM NaB for 72 hours were subjected to SDS_PAGE and western blot analysis. Monomeric involucrin was detected in all extracts and is indicated by the arrow. Cells were transfected with SVGFP or HPV16 and were co transfected with HIVtat. In cells transfected with SVGFP, a 130 kD band representing crosslinked involucrin was detected when cells are grown in the presence of NaB as indicated by the arrowhead. In cells transfected with HPV16 episomes, lower levels of the 130 kD band representing crosslinked involucrin were detected when cells are grown in the presence and absence of NaB as indicated by the arrowhead. Fold increase in crosslinked involucrin relative to monomeric involucrin is indicated in the graph. **E.** Cells were cotransfected with SVGFP or HPV16 and pCMVtat. Cell lysates from cotransfected OKF6tert1 cells treated with vehicle or 0. 0.25, 0.5 or 1 mM NaB for 72 hours were subjected to SDS_PAGE and western blot analysis. Levels of cytokeratin K10, loricrin, and acetylated H3 and HIVtat were detected in the extracts. Total H3 served as a loading control.

The effect of HIVtat on differentiation was examined in the presence of the HDAC inhibitor, NaB. Expression of involucrin was assessed and monomeric involucrin was expressed across conditions suggesting that at baseline these cells were differentiated. In cells expressing the SVGFP vector that were treated with NaB, an 8.6 fold increase was detected in a marker full terminal differentiation, crosslinked involucrin. This increase diminished to 5.9 fold with the co-expression of HIVtat. In SVGFP expressing cells, HIVtat alone did result in a 3.7 fold increase in crosslinked involucrin compared to vector alone, suggesting that HIVtat could facilitate further differentiation. In the context of HPV16 however, levels of cross linked involucrin were lowered substantially across all treatment conditions. In the presence of HPV16, HIVtat increased cross linked involucrin to the same levels as those attained with NaB treatment. HPV16 + NaB demonstrated a 1.4 fold increase compared to vector alone that was very similar to the 1.3 fold increase detected with HPV16 + HIVtat. HIVtat and NaB together resulted in an additive 2.4 fold increase in crosslinked involucrin in HPV expressing cells (Figure 2D).

Terminal and suprabasal differentiation markers were next assessed in the context of increasing NaB levels and of HIVtat (Figure 2E). Consistant with HDAC inhibition activity, increasing levels of AcH3 were detected with increasing concentrations of NaB (0, 0.25, 0.5 and 1mM) in SVGFP tranfected cells both in the presence and absence of HIVtat. Expression of HIVtat was confirmed by immunoblot (Figure 2E). Expression of HIVtat in the absence of NaB increased cell differentiation in SVGFP transfected cells as determined by increased expression of the terminal differentiation marker, loricrin, and decreased expression of the suprabasal marker, K10. This was consistent with the known inverse relationship that exists during normal epithelial differentiation that exists between expression of suprabasal markers such as K1 and K10 and terminal differentiation markers that include loricrin and fillagrin (21, 34). However in the presence of the differentiating agent, HIVtat decreased both Loricrin and K10 levels. In the presence of HPV16 and in the absence of NaB, HIVtat increased loricrin levels and did not change K10 levels. However, in the presence of HPV16 and increasing NaB levels, HIVtat increased both the suprabasal differentiation marker K10 and the terminal differentiation marker, loricrin (Figure 2E). Likewise in the presence of HPV16, HIVtat and NaB together demonstrated the highest levels of crosslinked involucrin (Figure 2D, lane 8). In the presence of HPV16 and a differentiating agent, HIVtat promoted both expression of suprabasal and terminal differentiation markers which may provide significant advantage to the HPV lifecycle (summarized in Table 1).

### HIVtat modulates late gene expression in HPV expressing cells induced to differentiate

Realtime qPCR was performed to assess later HPV events in the presence of HIVtat. The distribution of E6/E6* transcripts in the presence and absence of HIVtat and of NaB was assessed. The addition of HIVtat to HPV expressing cells, resulted in an increase in the spliced form of E6 (Figure 3A). A 7 to 8 fold increase in E6/E7 transcription was detected with the addition of NaB (Figure 3B). With NaB mediated full terminal differentiation, the spliced E6* form was consistently increased relative to unspliced E6/E7 (Figure 3B and 3C). With the addition of HIVtat the E6/E7:E6* ratios remained the same but the fold increase in transcription dimnished 2 fold overall (Figure 3B). Treatment of HPV expressing oral cells with HIVtat, NaB, or NaB + HIVtat demonstrated similar ratios of spliced to unspliced E6/E7 message (Figure 3C). A statitistically significant increase in HPV16 E1^E4 mRNA expression was detected in NaB treated HPV16/OKF6tert1 cells (p=0.02) (Figure 3D). Compared to to NaB treatment alone, NaB-induced OKF6tert1 cells that expressed HIVtat demonstrated a decrease in E1^E4 mRNA levels that was not statistically significant (Figure 3D). HIVtat alone did not increase E1^E4 levels in oral epithelial cells (Figure 3D) or in C33A cervical cells (Figure 3E). Interestingly, in the differentiated oral cells E1^E4 levels remained the same with the addition of HIVtat (Figure 3D left panel), but in the undifferetiated C33A cervical cells a statistically significant decrease in E1^E4 was detected in HIVtat expressing cells (p=0.01)(Figure 3D right panel). These experiments were performed three times, each time with triplicate experimental conditions. Late viral transcription events, E6* and E1^E4, including those induced by NaB diminished in the presence of HIVtat.

**Figure 3.**
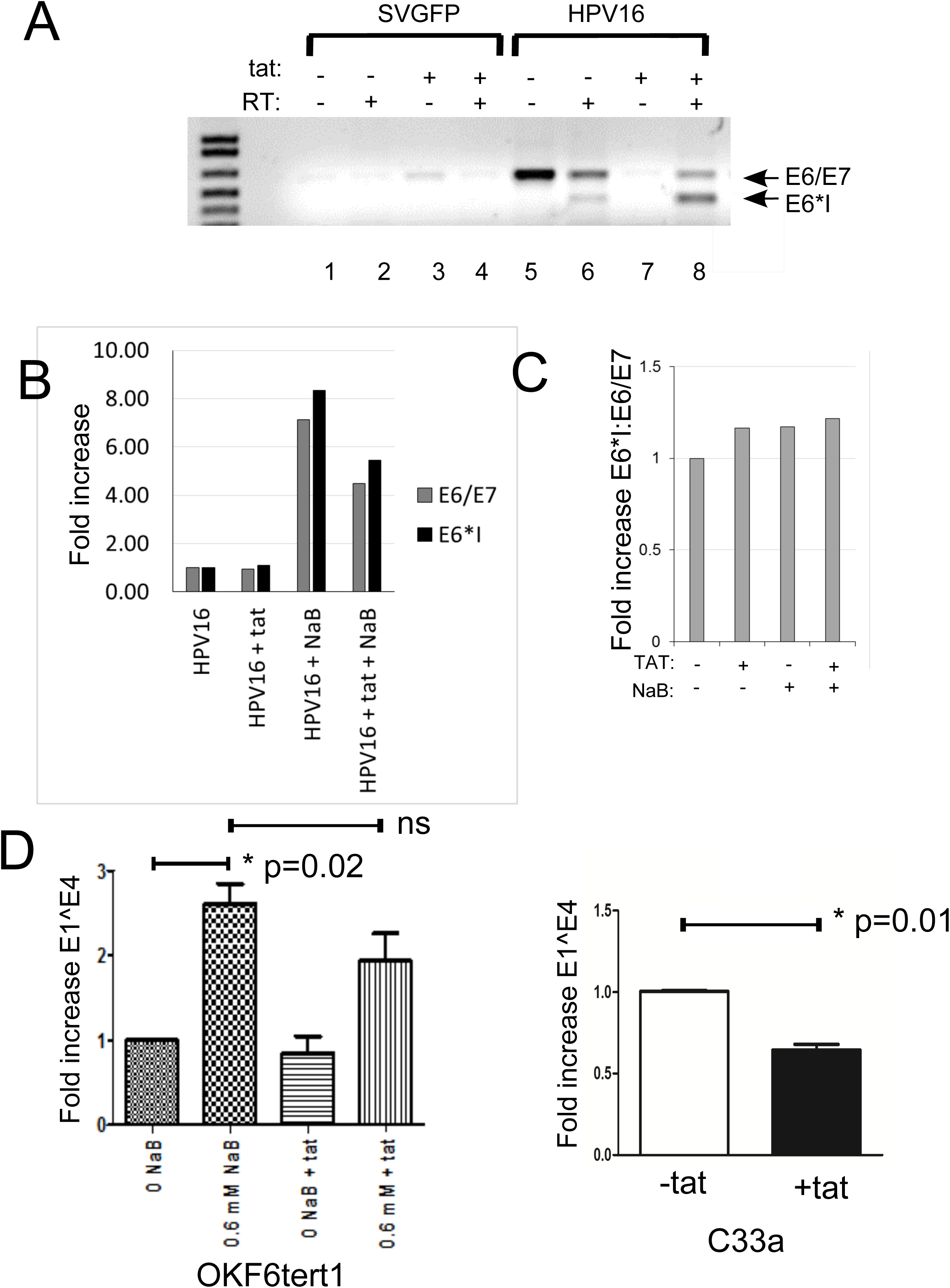
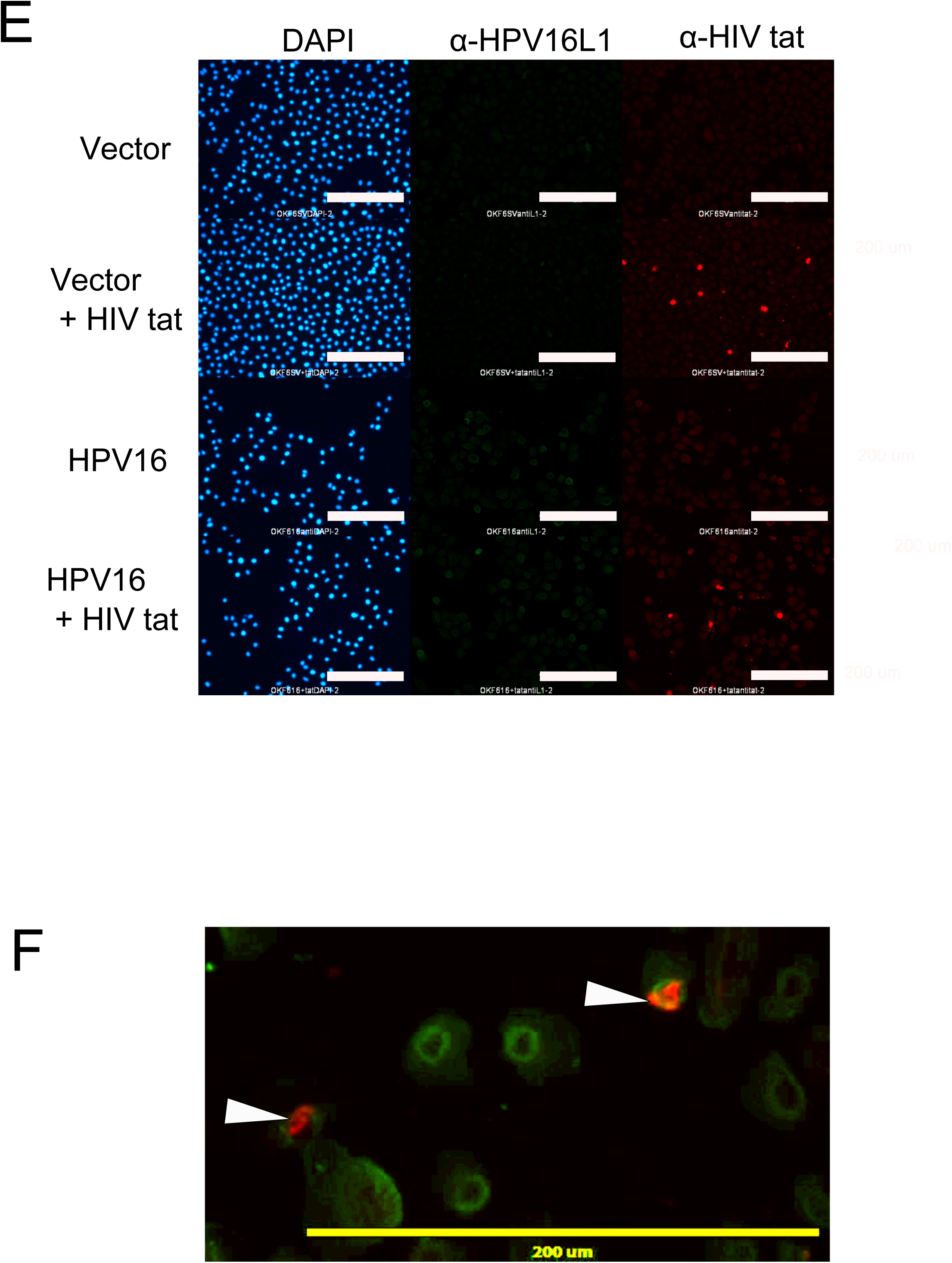
Treatment of OKF6tert1 cells with NaB or cotransfection with HIVtat results in HPV16 late gene expression. **A.** Following co transfection OKF6tert1 cells total RNA was isolated and used in RT-PCR to determine if HPV16 E6/E7 transcripts were present. The CaCl_2_ level in the media was raised to 2.0 mM to induce cellular differentiation. Total RNA was isolated from cells 48 hours after the increase in CaCl_2_ and used to make cDNA. E1^E4 levels were quantified by qPCR using primers that span the E1^E4 intron. Transcript levels were normalized to GAPDH. **B.** Following transfection OKF6tert1 cells with 0.6 mM NaB induce cellular differentiation. Total RNA was isolated from cells 48 hours after addition of NaB and used to make cDNA. E1^E4 levels were quantified by qPCR. Transcript levels were normalized to GAPDH. **C.** Reverse transcription reactions from transfected OKF6tert1 cells treated with NaB. The levels of spliced L1 transcripts were determined using L1 specific primers that span the intron. **D.** The levels of keratin K1 mRNA levels were determined using K1-specific primers. K1 mRNA expression was used to determine the extent of differentiation of treatment with NaB.

Potential interactions between HIVtat and HPV L1 were assessed. Perinuclear HPVL1 expression was detected in HPV/OKF6tert1 (HPV16L1 –Alexafluor 488 green) cells by immunofluoresence in the presence and absence of HIVtat (HIVtat Alexafluor 594-Bright red staining, Figure 3E). Confocal images demonstrate that HPV L1 (green) and HIVtat(red) were detected in a subset of the same cells (Figure 3F, white arrows). Transcripts containing L1 have repressive sequences that negatively regulate L1 expression (12, 52). The entire coding region of the HPV16 L1 gene (nt 5561-7156) was inserted directly 3’ to the GFP gene (Figure 4A, schematic). Reduced fluorecence was detected in OKF6tert1 cells expressing the pCMVGFP L1 construct, compared to cells expressing pCMVGFP L1 co-tranfected with HIVtat (Figure 4B, left panel). In the cotransfection, GFP expression levels were comparable to cells expressing GFP alone (Figure 4B, left and right panels). This suggested that inhibition of GFP expression by HPV16 L1 sequences was partially relieved by HIVtat (Figure 4B). Compared to GFP expressing cells, a 90% decrease in GFP mRNA was detected by qPCR in OKF6tert1 cells expressing GFPL1 in the absence of HIVtat (Figure 4B, right panel). In the presence of HIVtat, GFPL1 transcript levels were comparable to GFP alone (Figure 4B, right panel). The addition of HIVtat brought GFPL1 transcript levels back to baseline. To establish the region of L1 responsible for HIVtat-dependent restoration of the GFP signal, a series of overlapping deletions in the coding region of L1 were made within pCMVGFPL1 (Figure 4C, schematic left). Removal of the L1 region from nucleotide 5561 to 6820 (GFPL1E3) resulted in the the greatest production of GFP transcripts by qPCR (Figure 4C, right). Immunoblots also demonstrated that the greatest production of GFP protein was detected with the 5561 to 6820 (GFPL1E3) deletion mutant (Figure 4D, two biological replicates shown). While not statitstically signifcant, the GFPL1E5 appears to be HIVtat responsive. Both full length GFPL1 and GFPL1E5 demonstrate similar increases in GFP protein expression with HIVtat co-expression (Figure 4D). The presence of HIVtat increased GFP message and increased GFP protein expression 2 fold for deletion mutants spanning 5561-6820 (GFP5EL1). This suggested that L1 sequences within this region repressed GFP mRNA. Repressive sequences in this region of L1 have been previously described (28). Importantly, these results suggested that HIVtat may relieve this repression. In sum, HIVtat modulated late gene expression of E6*, E1^E4 and of L1 (summarized in Table 1).

**Figure 4.**
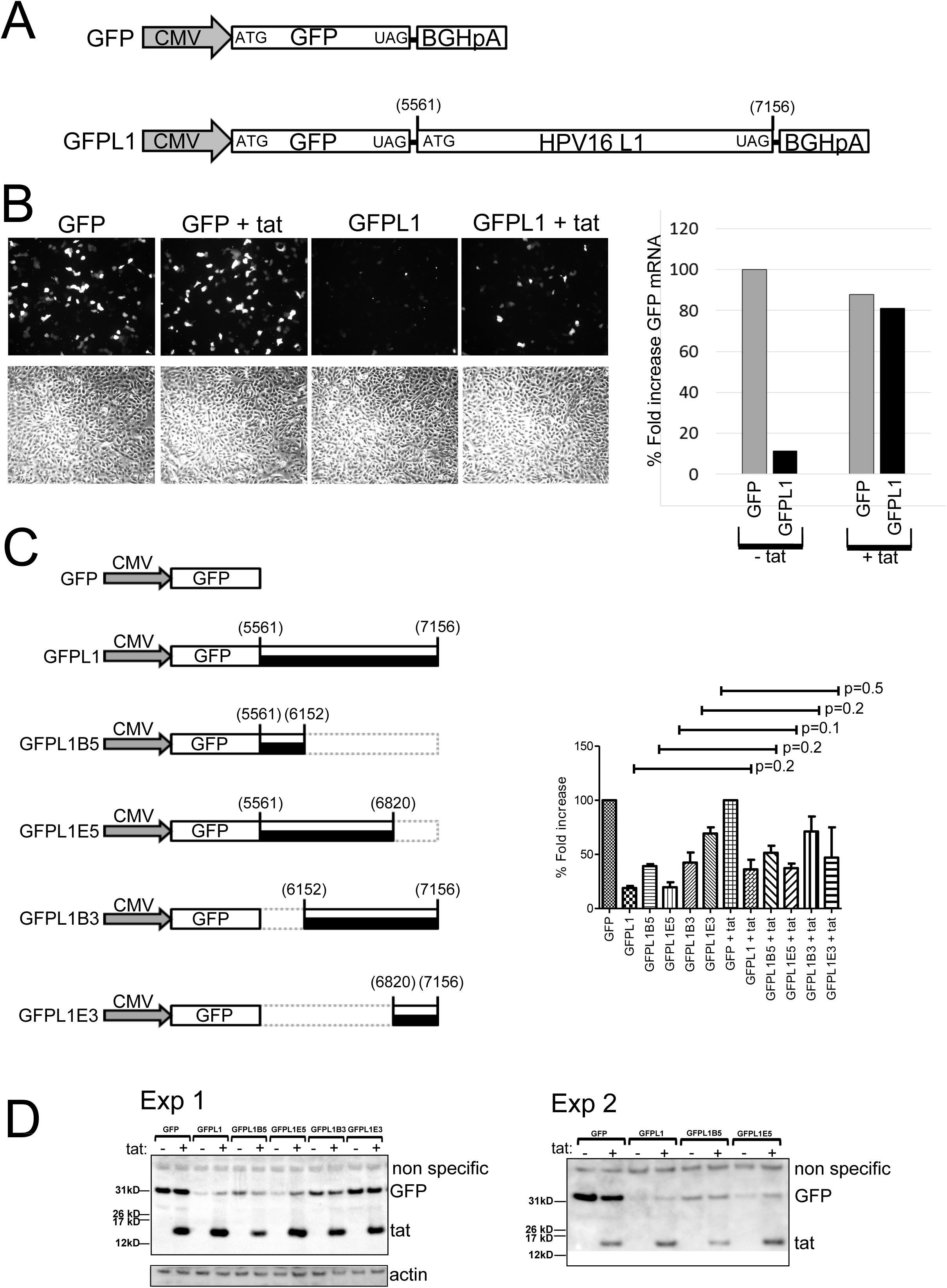
HIVtat stabilizes GFP transcripts containing L1 sequences. **A.** Schematic diagram showing the entire coding region of HPV16 L1 inserted 3’ to the coding region of GFP in pcDNA3GFP to produce pcDNA3GFPL1. The grey arrow indicates the CMV promoter driving expression in both constructs. The nucleotide positions of the HPV16 L1 start and stop codons are indicated. The bovine growth hormone polyadenylation signal is indicated by BGHpA. The start and stop codons of GFP are indicated by ATG and UAG, respectively. **B.** OKF6tert1 cells were cotransfected with either pcDNA3GFP or pcDNA3GFPL1 and either pCMV-HA or pCMV-tat. Cells were examined by light microscopy and GFP was monitored by fluorescent microscopy. RT-qPCR was performed on total RNA obtained from transfected cells to determine the effect of HIVtat on GFPL1 transcript levels and comparing them to GFP only levels. GFP-specific primers were used to quantitate GFP mRNA levels in the presence or absence of HIVtat. **C.** Serial deletions of HPV16 L1 DNA were made in pCMVGFPL1. The constructs were cotransfected into OKF6tert1 cells with or without HIVtat and GFP mRNA levels were detected by RT-qPCR. Deletion analysis localized the region of L1 necessary for HIVtat induced maintenance of L1 transcripts to a region spanning nucleotide 6152 to 6820. **D.** Western blot analysis of GFP protein levels. OKF6tert1 total cell lysates were obtained 48 hours after cotransfection of cells with GFPL1 deletion constructs and either control vector (pCMV-HA) or the tat expression vector, pCMV-tat. Western blot analysis was performed to determine GFP and HIVtat protein levels present in cell lysates. β-actin was detected to ensure equal loading of protein lysates. Exp. 1 and Exp.2 indicate western blot results from two separate experiments. **E.** HPV16 L1 and HIVtat protein expression was detected in OKF6tert1 cells by IFA. 48 hours post transfection, cells were fixed with methanol/acetone (1:1) and incubated with mouse α-HPV16L1 antibody (CAMVIR-1) and rabbit α-HIVtat antibody. Primary antibodies for HPV16 L1 and HIVtat were detected using alexafluor488 goat anti-mouse for (green) and alexafluor594 goat anti-rabbit antibody (red), respectively. Antibody staining was visualized by fluorescent microscopy. Cells were stained with DAPI to visualize nuclei. **F.** Coexpression of HPV16 L1 and HIVtat in individual cells. Following IFA signals for HPV16 L1 and HIVtat were merged to identify cells expressing both viral proteins.

To determine whether expression of HIVtat in HPV16/OKF6tert1 cells influenced production of viral particles, the level of HPV16 DNase-resistant DNA in the media of cells was determined. 10 ^5^ copies of HPV DNA were detected. HIVtat expression resulted in a 0.5 log increase in the levels of HPV16 particles detected in the culture media of HPV16/OKF6tert1 cells (Figure 5A). HIVtat associated 40nm HPV particles were morphologically identical to previously described native particles (Figure 5B, two representative virions are shown). In the presence of HIVtat and absence of NaB, an increase in copy number from 2.1×10^5^ mean copies/ml to 4.8 x10^5^ mean co/ml was detected (Figure 5A, Lanes 10 and 14). In the HIVtat+ NaB treated condition the copy number decreased from 2.1 x10^5^ mean co/ml to 1.2 x10^5^ mean co/ml (lanes 12 and 16). Media was collected from HIVtat expressing HPV16/ OKF6tert1 cells in the absence of NaB and used as an inoculum on naive OFK6tert1 cells. In differentiated cells infected with particles that were produced only in the presence of HIVtat, HPV16 E7 immunofluorescence was detected in over 50% of cells (Figure 5C). Supernatants from cells that did not express HIVtat were much less infectious (Figure 5C). Few cells were infected by particles generated in the absence of HIVtat.

**Figure 5.**
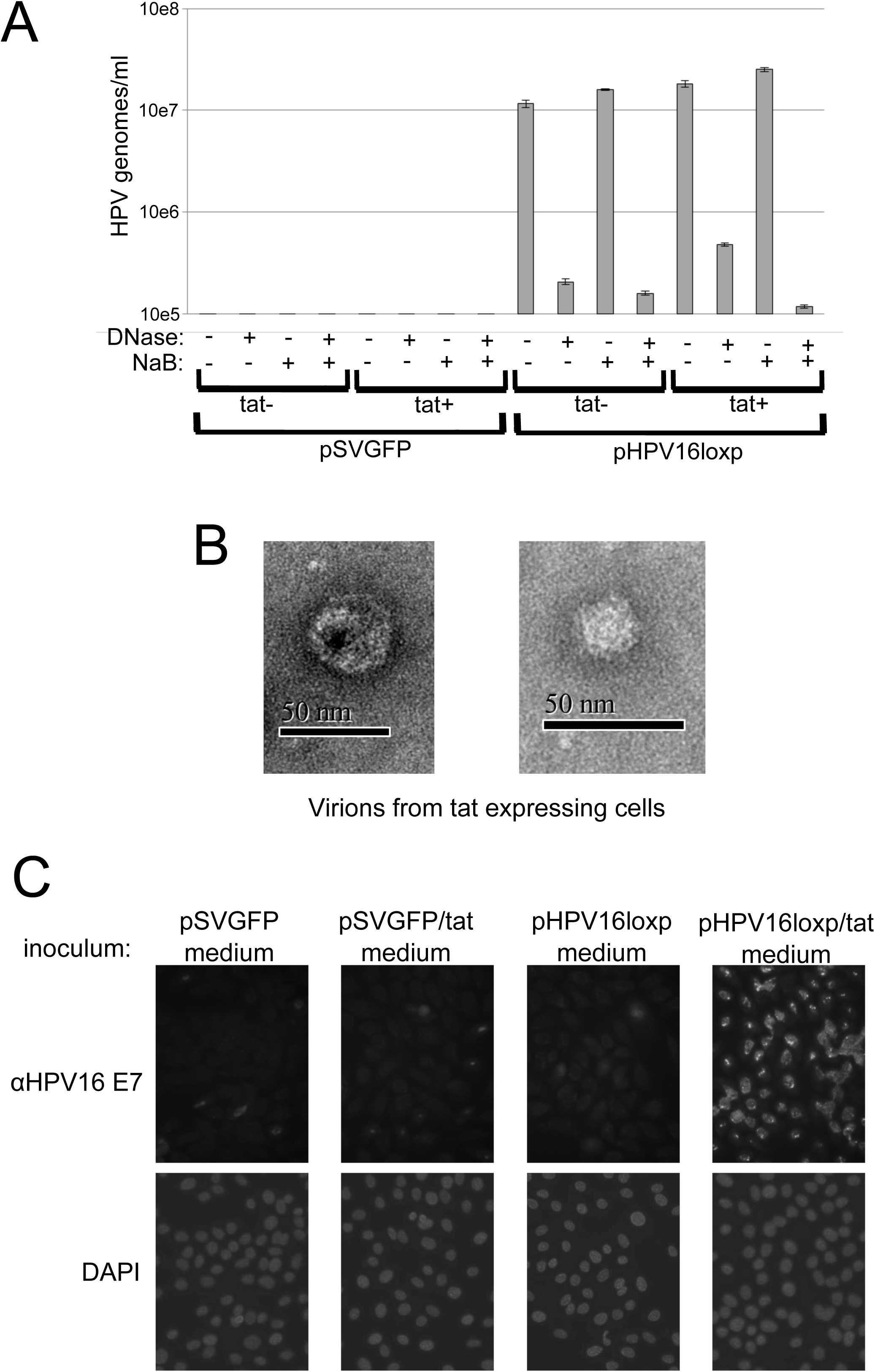
Successful de novo of infection of naïve OKF6tert1 cells by HPV16 requires expression of HIVtat in OKF6tert1 cells producing the inoculums. **A.** OKF6tert1 cells were cotransfected with either pSV40GFP (vector) or pHPV16loxp (HPV16) in the presence or absence of pCMV-tat (+/-tat). 24 hours post transfection, cells were treated with NaB (0.6 mM) in K-SFM without supplements (EGF/BPE) and incubated for an additional 72 hours. Media from cells was filtered and treated with DNase. DNA was isolated from filtered, DNase-treated media and HPV16 DNA was detected by qPCR using E7-specific primers. **B.** Filtered media from tat-expressing cells harboring HPV16 genomes was used to detect HPV16 particles by EM. **C.** Filtered media from OKF6tert1 cells cotransfected with either pSV40GFP (vector) or pHPV16loxp (HPV16) in the presence or absence of pCMV-tat (+/−HIVtat) were used to inoculate naïve OKF6tert1 cells. 48 hours post infection cells were fixed with methanol/acetone and stained with anti HPV16 E7 antibody. Nuclei were stained with DAPI following antibody incubation.

## Discussion

HPV is highly successful in the setting of HIV. It is well documented that HIV-infected individuals are at increased risk for the development of HPV-associated oral cancers compared to the general population (6, 11). HPV associated oral disease has persisted and increased during the era of highly active antiretroviral therapy (1, 10, 22, 36). While this could be associated with immune deregulation, the potential exists for direct virus-virus interaction. Little emphasis has been placed on the potential role of HIV-specific viral factors in the development of HPV-associated oral cancers and lesions. To our knowledge, the potential for direct HIV and HPV polyviral interactions in the oral cavity has not previously been explored. This paper describes the use of a simple cell culture system to assess the influence of HIVtat on HPV pathogenesis in oral keratinocytes.

HPV associated oral diseases are prevalent in the HIV positive population. There is a higher incidence of HPV related warts in this population (26) and these individuals shed HPV orally in up to 35% of cases (43). Importantly, the risk of HPV-associated oral cancers is 1.5-4.0 fold higher compared to the general population (6, 11). While compromised immunity surely plays a role in this increased risk it, does not fully explain the high rate of HPV-associated oral cancers (43). The biophysical properties of HIVtat make it a likely candidate for being involved in virus-virus interactions. There is evidence that HIVtat is secreted in individuals who are on antiretroviral therapy (32). HIVtat enhanced the pathogenesis of multiple opportunists including EBV (13), KSHV (53) and BKV (24).

The study of oral HIVtat and HPV interactions are feasible. HIVtat is produced in and secreted from HIV infected cells and has been detected in the saliva of HIV infected individuals (48). It has previously been shown that HIVtat can increase the proliferative potential of HPV positive oral keratinocytes. This was demonstrated by HIVtat mediated increased expression of viral oncogenes E6/E7 and increased cellular proliferation in vitro in the presence of serum and calcium (29). HIVtat also facilitated HPV E6/E7 associated nodule formation, *in vivo* in nude mice (29). Secreted HIVtat is readily transduced into uninfected cells and, therefore, it can exert its regulator effects on a cell that does not produce it (8). These effects included HIVtat’s ability to regulate cellular gene expression, induce DNA damage, and change the redox potential of the cells (4, 9, 51). In this report, HIVtat induced a DNA damage response in oral keratinocytes, and demonstrated a modest effect on the expression of cellular genes involved in sensing oxidative stress. Interestingly, HIVtat’s transactivation potential, modulation of oxidative stress, and mediated DNA damage did not enhance HPV pathogenesis in oral epithelial cells (Figure 2).

HIVtat, however, did modulate oral epithelial cell differentiation. A hallmark of epithelial differentiation, is exit from the cell cycle. In the context of HPV, HIVtat has been previously shown to promote cell cycle progression, through decreased expression of cell cycle inhibitors (35). We demonstrate that in the presence of HPV16 and the differentiating agent, NaB, HIVtat promoted expression of both the suprabasal marker, K10, and the terminal differentiation marker, loricrin (Figure 2B). NaB induced full terminal differentiation as demonstrated by increased expression of crosslinked involucrin. Full terminal differentiation, as determined by detection of cross linked involucrin, was not as robust as in the absence of HPV. Both in the presence and absence of HPV, HIVtat increased full terminal differentiation as determined by crosslinked involucrin levels. Interestingly, across the experimental conditions, HPV decreased expression of crosslinked involucrin. In the absence of HPV, HIVtat decreased NaB induced cross-linked involucrin levels fourfold (Figure 2A). In sum, HIVtat and HPV16 together promote, a cellular phenotype with enhanced suprabasal and terminal differentiation features that does not reflect full terminal differentiation.

These differentiation associated cellular responses induced by HIVtat have been shown to be important for the normal HPV life cycle and suggested that introduction of HIVtat into an HPV16-infected oral keratinocyte may impact viral production in this cell type. Taken together, the above data suggest that HIVtat and HPV collectively produce an enhanced suprabasal state, important to early gene expression, and early terminal differentiation state, important to viral amplification and late gene expression, that could be significantly advantageous to the HPV lifecycle. HIVtat expression resulted in enhanced production of E6* transcripts (Figure 3A) and levels of diminshed E1^E4 (Figure 3D) in both oral and cervical epithelial cells. Compared to the inducer of terminal differentiation alone, the presence of both NaB and HIVtat consistently performed poorer as a promoter of HPV gene expression (Figure 3). E1^E4, is important to cytokeratin destabilitation and to viral egress. HIVtat has been shown by others to destabilize the cytoskeleton (3, 31). Further in a recent study of the oral epithelial proteome of HIV infected subjects, upregulation of the breakdown products associated with the intermediate fillament, vimentin were detected (50). This also suggests destabilization of the cytoskeleton by HIV. These processes may facilate HPV egress in the absence of E1^E4.

Properly spliced L1 is critical to virus production. In the study presented here, HIVtat was detected in a subset of L1 expressing oral epithelial cells (Figure 3E). Perhaps, HIVtat directly influenced L1 production. Previous studies have demonstrated repressive sequences within the L1 open reading frame. Overexpression of heterogeneous nuclear ribonucleoprotein (hnRNP) has previously been shown to induce HPV16 late gene expression from HPV16 episomal genomes. Binding of hnRNP C1 to the HPV16 early, untranslated region activated a HPV16 late 5’-splice site that resulted in production of HPV16 L1 mRNAs (17). Here we show that HIVtat relieved L1 imposed repression of transcription that mapped to nucleotides 5561-6820, a known inhibitory region (Figure 4). It has previously been shown in cells harboring integrated HPV 18, that L1 transcripts were increased upon HIVtat treatment (19). The HIVtat mediated modification of oral epithelial differentiation (shown in Figure 2) may modulate the cellular factors like the differentiation associated hnRNPs that contribute to splicing of both cellular and viral genes. Importantly, hnRNPs that are critical to splicing have recently been shown to cooperate with RNA helicases to regulate cellular differentiation (16). Future studies will need to be performed to determine potential that HIVtat is able to exploit this process.

HIVtat expression allowed for a modest increase in virion production (summarized in Table 1; Figure 6). In the presence of HIVtat, the L1 protein native to the clinical HPV isolate was generated and may have allowed for productive infection. It could be that the modest HIVtat effect on oxidative stress genes resulted in an oxidative state favorable to L1 conformation necessary for infectivity (14). In other systems, L1 has been provided in trans to facilitate particle production (40). We have demonstrated for the first time that the addition of HIVtat to our episomal HPV system resulted in the production of HPV16 particles with increased infectivity compared to virus produced in HIVtat negative cells (Figure 5). HIVtat appeared to work with HPV to generate an optimal suprabasal and early terminal differentiation phenotype that may be conducive to HPV replication in the context of HIV. HIVtat associated HPV particles were more infectious than those generated in the absence of HIVtat. Given HIVtat’s ability to be easily taken up by multiple cells, we can also speculate that HIVtat may act as a chaperone, ushering HPV virions into naive cells (Figure 6B). The potential for this observation will need to be addressed in future studies. In sum, differentiated oral epithelial cells support the HPV life cycle in the presence and absence of HIVtat (Figure 6A and B), however, HIVtat modified the differentiation state and resulted in more successful de novo infection.

**Figure 6.**
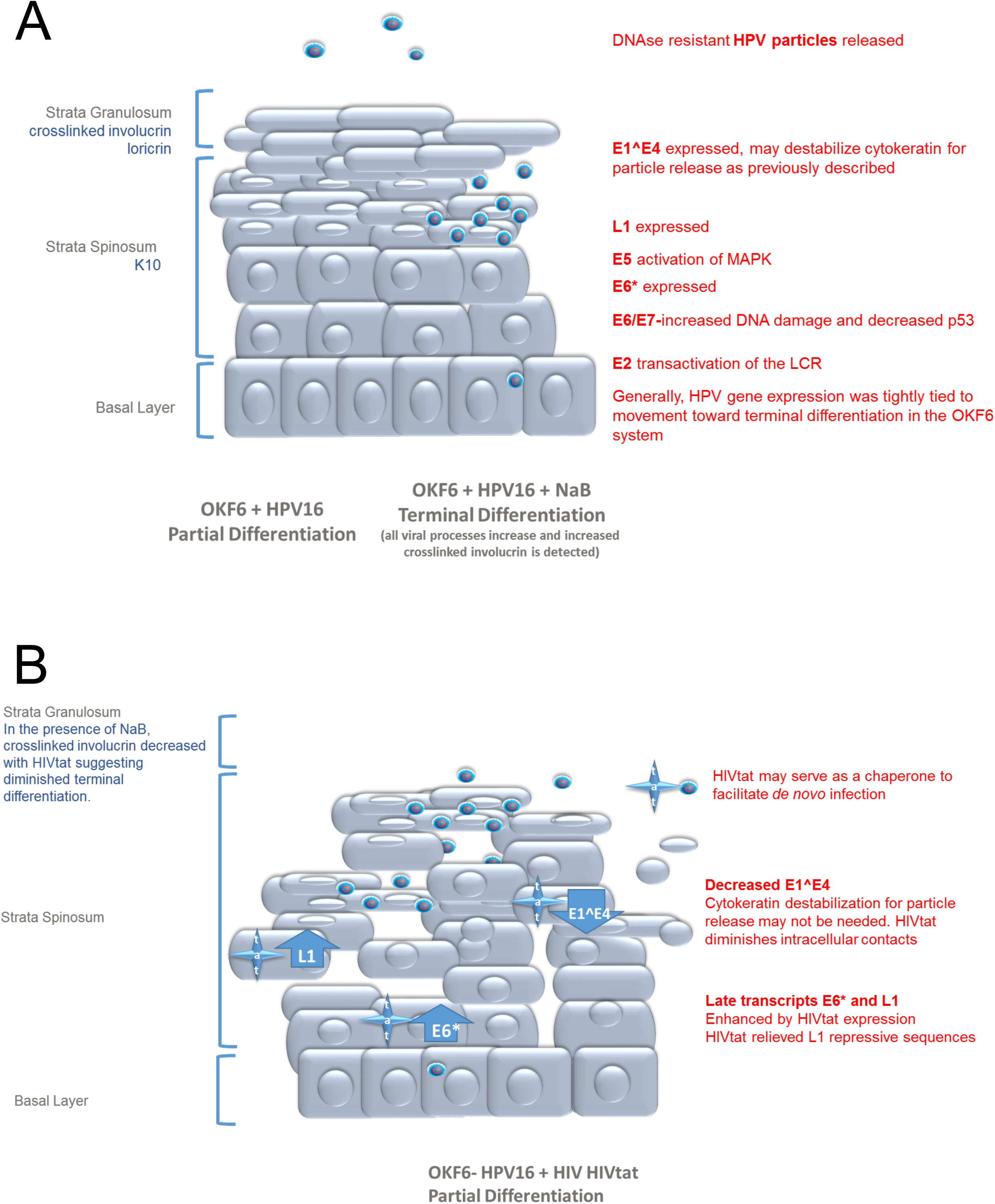
Effect of HIVtat on HPV16 gene expression and virus production. **A.** graphical representation OKF6 cells and associated differentiation markers. **B.** A graphical representation HPV infected oral epithelial cells in the presence and absence of HIVtat. Viral amplification and release shown within the oral epithelium (center). Expressed epithelial differentiation markers are shown adjacent to epithelial layers (left). HIVtat modulation of HPV gene expression and life cycle related cellular processes are shown adjacent to epithelial layers.

The above observations of direct HPV/HIV interactions could result in significant impact in a clinical setting regarding the acceleration of HPV in the context of HIV. The potential for immune reconstitution-associtated promotion of HPV in the setting of HIV has recently been described (43). Potential consequences of these direct and indirect interactions could manifest as enhanced oncogenesis and increased spread of HPV-associated disease. Theoretically, theraputic interventions focused on targeting of HIVtat could be beneficial to decreasing HPV-associated disease. This investigation provides evidence that HIVtat may contribute to the increased HPV16-associated oral cancers detected in conjunction with HIV infection. This mechanism may also drive the many other HPV types that are associated with oral warts.

## Acknowledgements

We would like to thank James G. Rheinwald for providing us with the OKF6tert1 cells used in this study and Marian Couch and Xiao Ying Yin for providing us with oral cancer biopsy DNA used to make whole HPV genomes. This research was supported by NCI supplemental funding for HIV-associated malignancy research to the UNC Lineberger Cancer Center and the UNC CFAR (P30-CA016086), NIH/NIDCR and NIAID (OHARA) 1U01AI068636 and NIH/NIDCR 1R56DE023940-01.

---

**Table 1. OKF6tert1 cells are differentiated and support HPV viral replication**. **A.** Summary of OKF6tert1 cells differentiation state with and without NaB and HIVtat. **B.** Visual and graphical summary of OKF6tert1 cells containing HPV16 episomes. Differentiation state and viral gene expression were examined with and without NaB and HIVtat. (−) not detected (+) detected (ND) not done

