## Supplemental Table 1 for "HIVtat Alters Epithelial Differentiation State and Increases HPV16 Infectivity in Oral Keratinocytes"

Supplemental Table 1. Sequence of oligonucleotide primers used for PCR and qPCR

| Primer | Sequence |
| --- | --- |
| HPV16LCRREPF | 5’GCGACGCGTGACCTAGATCAGTTTCCTTTAGGACG3’ |
| HPV16LCRREPR | 5’GCGAGATCTATAAAATGTCTGCTTTTATACTAACCGGT3’ |
| HPV16E6EXF | 5’ CGCAAGCTTGGATGCACCAAAAGAGAACTGCA3’ |
| HPV16E6EXR | 5’ CGCGAATTCTTACAGCTGGGTTTCTCTACG3’ |
| HPV16E7EXF | 5’CGCAAGCTTATGCATGGAGATACACCTACATTG3’ |
| HPV16E7EXR | 5’ CGCGAATTCTTATGGTTTCTGAGAACAGATGGG3’ |
| HPV16E2EXPF | 5’ GGGAAGCTTGCCATGGAGACTCTTTGCCAACGTTT3’ |
| HPV16E2EXPR | 5’GGGACTAGTTCATATAGACATAAATCCAGTAGACACTGT3’ |
| HPV16E1^E4F | 5’CCCCATCTGTTCTCAGAAACC3’ |
| HPV16E1^E4R | 5’GGCCAATGTCTGCCTAATAA3’ |
| HIVTATFF | 5’GCGCATATGGAGCCAGTAGATCCTAGA3’ |
| HIVTAT3’R | 5’CCCGAATTCCTATTCCTTCGGGCCTGTCGGGT3’ |
| HumanGPX1F | 5’CCCTCTCTTCGCCTTCCT3’ |
| HumanGPX1R | 5’CAACATCGTTGCGACACAC3’ |
| HumanSOD2F | 5’GCAAGGAACAACAGGCCTTA3’ |
| HumanSOD2R | 5’GTAGTAAGCGTGCTCCCACAC3’ |
| LUCF | 5’CCAAAATTTACATTAGGAAAACGAA3’ |
| LUCR | 5’TTTATGTTGCATGACACAATAGTT3’ |
| GFPqPCRF | 5’GAAGCGCGATCACATGGT3’ |
| GFPqPCRR | 5’CCATGCCGAGAGTGATCC3’ |
| GAPDHF | 5’GAAGGTGAAGGTCGTAGTC3’ |
| GAPDHR | 5’GAAGATGGTGATGGGATTTC3’ |
